# Transcriptome data from human nasal epithelial cells infected by H3N2 influenza virus indicate early unbalanced ROS/RNA levels, temporarily increased aerobic fermentation linked to enhanced α-tubulin and rapid energy-dependent IRF9-marked immunization

**DOI:** 10.1101/2021.10.18.464828

**Authors:** José Hélio Costa, Shahid Aziz, Carlos Noceda, Birgit Arnholdt-Schmitt

## Abstract

**Background:** Transcriptome studies of a selected gene set (*ReprogVirus*) had identified unbalanced ROS/RNS levels, which connected to increased aerobic fermentation that linked to alpha-tubulin-based cell restructuration and cell cycle control, as a *ma*jor *c*omplex *t*rait for *e*arly *d*e novo programming (*CoV-MAC-TED*) upon SARS-CoV-2 infection. Recently, CoV-MAC-TED was confirmed as promising marker by using primary target human nasal epithelial cells (NECs) infected by two SARS-CoV-2 variants with different effects on disease severity. To further explore this marker/cell system as a standardized tool for identifying anti-viral targets in general, testing of further virus types is required. Results: Transcriptome level profiles of H3N2 influenza-infected NECs indicated ROS/RNS level changes and increased transcript accumulation of genes related to glycolysis, lactic fermentation and α-tubulin at 8 hours post infection. These early changes linked to energy-dependent, IRF9-marked rapid immunization. However, *ReprogVirus*-marker genes indicated the absence of initial cell cycle progress, which contrasted our findings during infections with two SARS-CoV-2 variants, where cell cycle progress was linked to delayed IRF9 response. Our results point to the possibility of CoV-MAC-TED-assisted, rapid individual host cell response identification upon virus infections. Conclusion: The complex trait CoV-MAC-TED can identify similar and differential early responses of SARS-CoV-2 and influenza H3N2 viruses. This indicates its appropriateness to search for anti-viral targets in view of therapeutic design strategies. For standardization, human NECs can be used. This marker/cell system is promising to identify differential early cell responses upon viral infections also depending on cell origins.

## 1. Background

Influenza A subtype H3N2 is a respiratory single-stranded RNA virus. It causes worldwide seasonal diseases in the upper respiratory tract. Human nasal epithelial cells (NECs) are the primary targets for influenza infections. Host response in these cells is critical in general for resultant pathologies of respiratory viruses including also coronaviruses. Therefore, a group of clinical researchers from Singapore established a powerful in vitro system of cultured NECs that enables deeper insights into early host cell responses (1, 2). We took profit of public RNA sequencing data from this group to test our recently published approach and identify traces of the complex marker system ‘CoV-MAC-TED’ during early host responses at 8, 24 and 48 hours post infection (hpi) by influenza H3N2 strain.

CoV-MAC-TED refers to *ma*jor *c*omplex *t*rait for *e*arly *d*e novo programming and was developed from a mixed approach of plant and coronavirus research (3). It is based on a set of genes, named *ReprogVirus* (4). Recently, we successfully detected CoV-MAC-TED by help of a selected core subset of *ReprogVirus* genes in the described human NECs infected by SARS-CoV-2 variants (5). We also pointed to the lack of recognizing early metabolic-structural reorganization as an essential part of immunology across organism boarders (4, 5).

The core subset of genes characterizing CoV-MAC-TED was selected to identify a shift in ROS/RNS (ASMTL, SOD1, SOD2, ADH5, NOS2), to represent glycolysis (HK, PFK, GAPDH, Eno) and lactic acid fermentation (LDH) as well as structural cell organization (α-Tub). We assumed that ASMTL could indicate oxidative stress equilibration, while SOD1 and SOD2 genes mark anti-oxidative activities and were selected to indicate oxidative stress. ASMTL is a paralog of ASMT, which is involved in melatonin synthesis in human cells (7). However, we could not find ASMT gene transcripts in collected human NECs. ADH5 is known to be involved in ROS/RNS equilibration through NO homeostasis regulation (8, 9, 10) and the inducible NOS gene, NOS2, relates to the induction of NO production (6, 11). NOS1 and NOS3 had not been encountered in sufficient quantities in the collected epithelial nose cell data. Further, we selected SNRK and mTOR to highlight cell energy-status signaling (11, 12, 13, 14, 15). The mTOR is activated when there is excess of energy in contrast to SNRK, where higher expression indicates energy depletion. Genes for E2F1 and mTOR were included to indicate changes in cell cycle regulation, namely cell cycle progress (G1/S and G2/M transitions) (15). E2F1 belongs to the transcription factor family E2F and is known as cell cycle activator. The interferon regulator factor, IRF9, demonstrated early transcription in SARS-CoV-2 infected human lung adenocarcinoma cells and was therefore proposed as functional marker candidate that could signal initiation of the classical immune system (3).

Viruses can affect host cell cycle regulation in favor of viral replication, which might seriously affect host cell physiology with impacts on pathogenesis (16–19). Influenza A virus was found to arrest cell cycle in in G0/G1 Phase (20, 21). Corona viruses have been shown to notably arrest cells in G0/G1 stage of cell cycle (18) identified 12h, 24h, and 48h signatures from calu-3 lung adenocarcinoma cells infected with SARS-CoV, which included genes related to cell cycle progress, viz., E2F and mTOR. Enrichment of coronavirus-infected cells had also been found in G2/M stage (22).

## 2. Material and Methods

### 2.1 Gene expression analyses of transcriptomic data from H3N2 influenza infected human nasal epithelial cells

The gene expression analyses of the main ReprogVirus genes (3, 4) were performed in transcriptomic data from human NECs infected with H3N2 influenza virus at times 0, 8, 24 and 48 hpi (hours post infection). The RNA-seq data, previously deposited in GenBank (NCBI) by Tan et al. (1), are available in SRA database under the bioproject PRJNA566434. The studied ReprogVirus genes were: ASMT, ASMTL, SOD1, SOD2, ADH5, NOS1, NOS2, NOS3, Total HK (HK1, HK2, HK3), Total PFK (PFKM, PFKL and PFKP), GAPDH, Total Eno (Eno1, Eno2 and Eno3), Total LDH (LDH-A, LDH-B, LDH-C, LDH-AL6A, LDH-AL6B), Total α-Tub (TUB-A1B, TUB-A1C, TUB-A4A), SNRK, mTOR, E2F1 and IRF9. The accession numbers of these genes are available in Costa et al. (3). In addition, transcript levels of H3N2 influenza proteins as polymerase PB2 (AHL89269.1), polymerase PB1 (YP_308847.1), polymerase PA (YP_308846.1), hemagglutinin (YP_308839.1), nucleocapsid protein (YP_308843.1), neuraminidase (YP_308842.1), matrix protein 1 (YP_308841.1), nonstructural protein 1 (ABB04321.1) were evaluated to monitor the virus proliferation. The gene expression analysis was performed as described in Costa et al. (5) in this Special Issue.

### 2.2 Statistical analyses

Normality and homogeneity of variances from the analyzed variables (in RPKM) were tested with Shapiro-Wilk test and Levene tests, respectively, using InfoStat 2018I. Only in one case (SOD2) data had to be converted to parametric for further analyses (applying squared root). Then, ANOVA tests for repetitions along time in the same donor were performed by using Excel datasheet. Significance levels were set at α=0.01 and α=0.05.

Regression and correlation tests were performed to evaluate possible associations between different cell transcript increases or levels regarding cell donors. Also, different virus transcripts were submitted to regression and correlation analyses against IRF9 transcripts. Several types of regressions were considered with Excel software. Correlations with p<0.05 were considered significant.

We highlight that we interpret our data as ‘real’ observations under the employed conditions involving only small samples, which certainly provide insights that cannot get relevance or not relevance by calculating significance and performing correlation analyses. Nevertheless, we applied these calculations at usual p-values for biological research as an aid to focus our insights. Readers are encouraged to making themselves familiar with the current paradigm change related to the usage of statistical significance (23, 24, 25, 26, and 27).

## 3. Results

In **Figure 1**, transcript accumulation of selected ReprogVirus genes at 8 h, 24 h and 48 h post infection (hpi) with influenza H3N2 is shown in % of 0 hpi for an average of seven cell donor origins. RPKM values can be consulted in **table S1 (Supplementary File S1)**.

**Figure 1:**
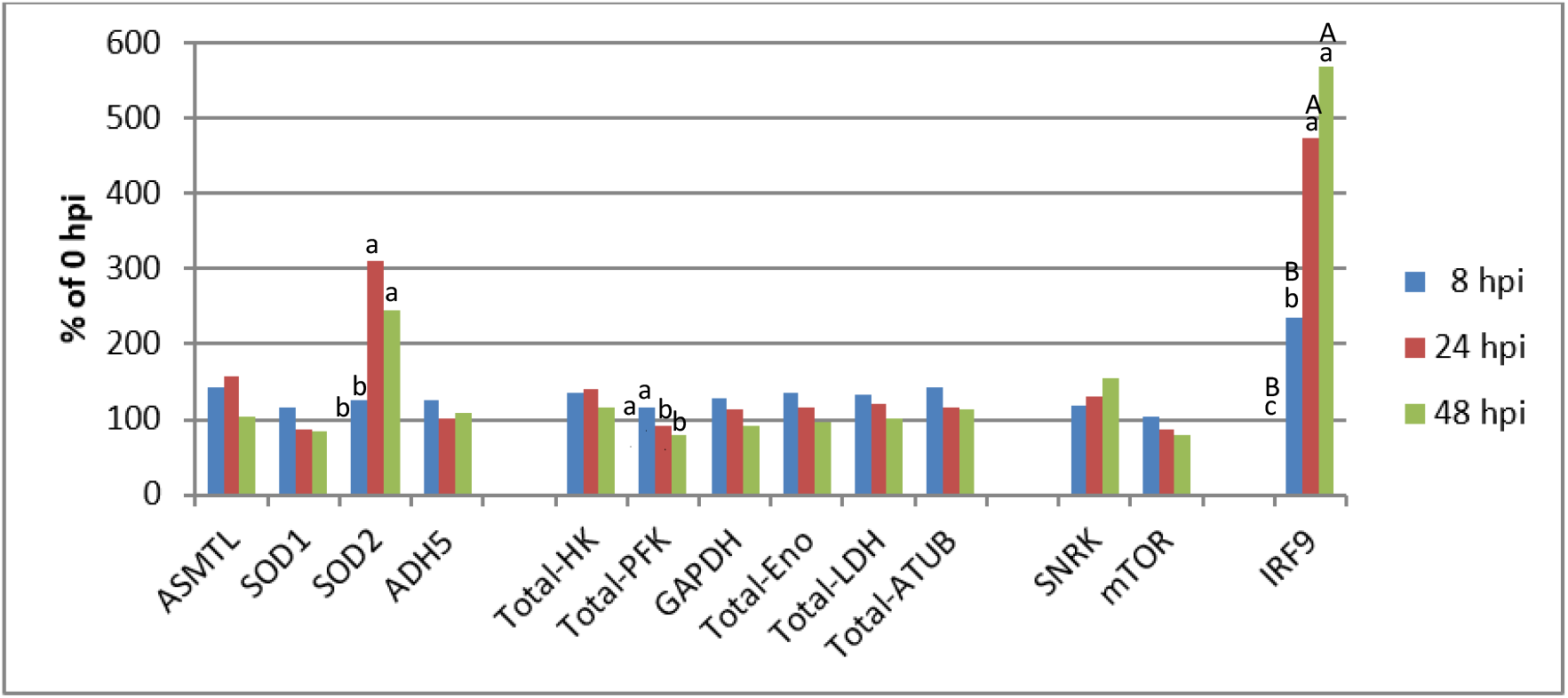
Early transcript accumulation of selected *ReprogVirus* marker genes in human nasal epithelial cells infected with H3N2 Influenza virus. These data were obtained from 7 different cell origins according Tan et al. (1). The RPKM average values can be found in supplementary table S1. Different letters indicate significant differences between net RPKM: upper and lower case for α=0.01 and 0.05, respectively. Letters on the 100% horizontal line correspond to 0 hpi.

ASMTL and ADH5 transcript levels signal rapidly increased oxidative stress at 8 hpi related to ROS and RNS. Also, SOD1 and SOD2 (significant) indicate this early oxidative stress by ROS, which confirms functionality of our marker gene candidate ASMTL. None of the three NOS genes was found sufficiently expressed for data integration into this study, and, in fact, ADH5 signaled at 8 hpi only a moderate increase in transcription of 125% compared to 144% by ASMTL. However, more clearly unbalanced ROS/RNS was signaled for 24 hpi. ASMTL reached a level of 157% against basal levels observed for ADH5. This signaling of ASMTL is supported by highly increased transcription for SOD2 to a level of 311% (significant), which again confirms credibility of our target marker system ASMTL/ADH5 to mark ROS/RNS unbalancing. This early increase in ROS-related stress associates to a moderate increase in transcript levels for genes related to glycolysis (115% to 140% depending on the gene and time) at 8 hpi and 24 hpi, but a significant decrease for PFK was noticed from 8 to 24 hpi. This linked to similar increase in LDH transcription of 133% at 8 hpi with slightly lower levels of 121% at 24 hpi. Rapid increase of 143% for α-tubulin transcript accumulation at 8 hpi but no indication for cell cycle progress through low levels for E2F1 and basal levels for mTOR point to rapid cell reorganization that did not include a change in cell cycle regulation. Transcript levels for both α-tubulin and mTOR were decreased again to basal respectively below basal levels from 24 hpi. However, SNRK signaled energy depletion at 8 hpi (118%), which increased steadily from 24 hpi (131%) to 48 hpi (156%). This increase in SNRK-signaled energy depletion accompanied strong enhancement of transcript levels for IRF9, which is expected to mark the formation of the overall classical immune system. IRF9 demonstrated early significant (234% at 8 hpi) and highly significant (473% at 24 hpi) increase and reached the highest enhancement in transcript accumulation at the finally included time point 48 hpi (568%). This last observation confirms the quality of IRF9 as a marker candidate for the classical immune system.

Correlation analyses between transcript levels or increases of gene pairs at different times post infections underlined the relevance of early changes in oxidative stress (ROS) for adaptive host cell energy management (glycolysis and aerobic fermentation) and its link to the classical immune system marked by IRF9. Throughout the seven cell donor origins, we found that levels of SOD2 transcripts at 8 hpi were linear and positively correlated (p<0.05) with those of total PFK, total Eno, total LDH and SNRK at 24 hpi (**Supplementary File S2, Table S2**). Also, there was significant quadratic correlation (R^2^ = 0.83) between increase in transcript levels of SOD2 from 0 to 24 hpi and ADH5 at 24 hpi. This correlation manifested as decrease or increase of ADH5 transcript levels at low or high increases of SOD2 transcript levels (**Supplementary File S2, Figure S1**). Furthermore, the fold increase in total PFK transcripts at 8 hpi was positively correlated with that of IRF9 from 8 to 24 hpi. Transcript levels of SOD1 and of ASMTL at 24 hpi correlated positively with fold increase of IRF9 transcripts from 0 to 24 hpi. To the contrary, rapid fold increase in IRF9 transcript levels early from 0 to 8 hpi was negatively correlated (p<0.05) at linear regression to fold increase of total PFK and Eno levels from 8 to 24 hpi along with those of total α-Tub, mTOR, SOD1 and ADH5 (**Supplementary File S3**).

**Figure 2** shows early transcript accumulation of selected *ReprogVirus* genes in human NECs of individual profiles of the seven donor origins that are underlying the averaged analysis displayed in Figure 1. In **Figure 2A**, the transcript accumulation profiles at 0 hpi are presented for RPKM, whereas **Figures 2B – 2D** indicate the change in transcript levels at 8 hpi (Figure 2B), 24 hpi (Figure 2C) and 48 hpi (Figure 2D) in % of 0 hpi. It can be seen in Figure 2A that transcript levels of the selected genes in RPKM mirror high individual diversity for genes related to glycolysis, LDH and α-tubulin. To our best knowledge, the studied cell cultures were not synchronized before virus infection (1, 2). So, it is not possible to judge whether this variability is due to cell origin from a donator or whether also cell culture conditions and shifted cell cycle phases or all together were causing this variability. A weakness comes also through the fact that only one technical repetition was available for RNA sequencing data. Nevertheless, preliminary analysis was performed and indicated meaningful individual diversity of responses upon virus infection. This variability is reduced among the transcripts for each gene when the change in transcript levels upon virus infection is considered in % of 0 hpi. This way of data presentation enabled highlighting individual cell origins that respond in an exceptional way. E.g. D02 at 8 hpi shows the highest ASMTL-marked ROS stress associated to rapidly unbalanced ROS/RNS, which links to the earliest immune response already at 8 hpi, if marked by IRF9. On the contrary, for D05 ADH5 transcript levels indicate early and over time highest NO stress among all seven origins, leading to unbalanced ROS/RNS in favor of RNS. It links to the highest immune response from 24 hpi. It also links to the highest initial increase in transcript levels for glycolysis (GAPDH and Eno), LDH, α-tub, SNRK and mTOR with striking increase of the energy-depletion indicator SNRK at 24 hpi and 48 hpi accompanied by enhanced IRF9 response. Furthermore, for D01 a continuously balanced relationship is indicated from 8 hpi between ROS and RNS marked by ASMTL and ADH5 that associates to the lowest relative IRF9 response over all time points among the seven cell origins.

**Figure 2:**
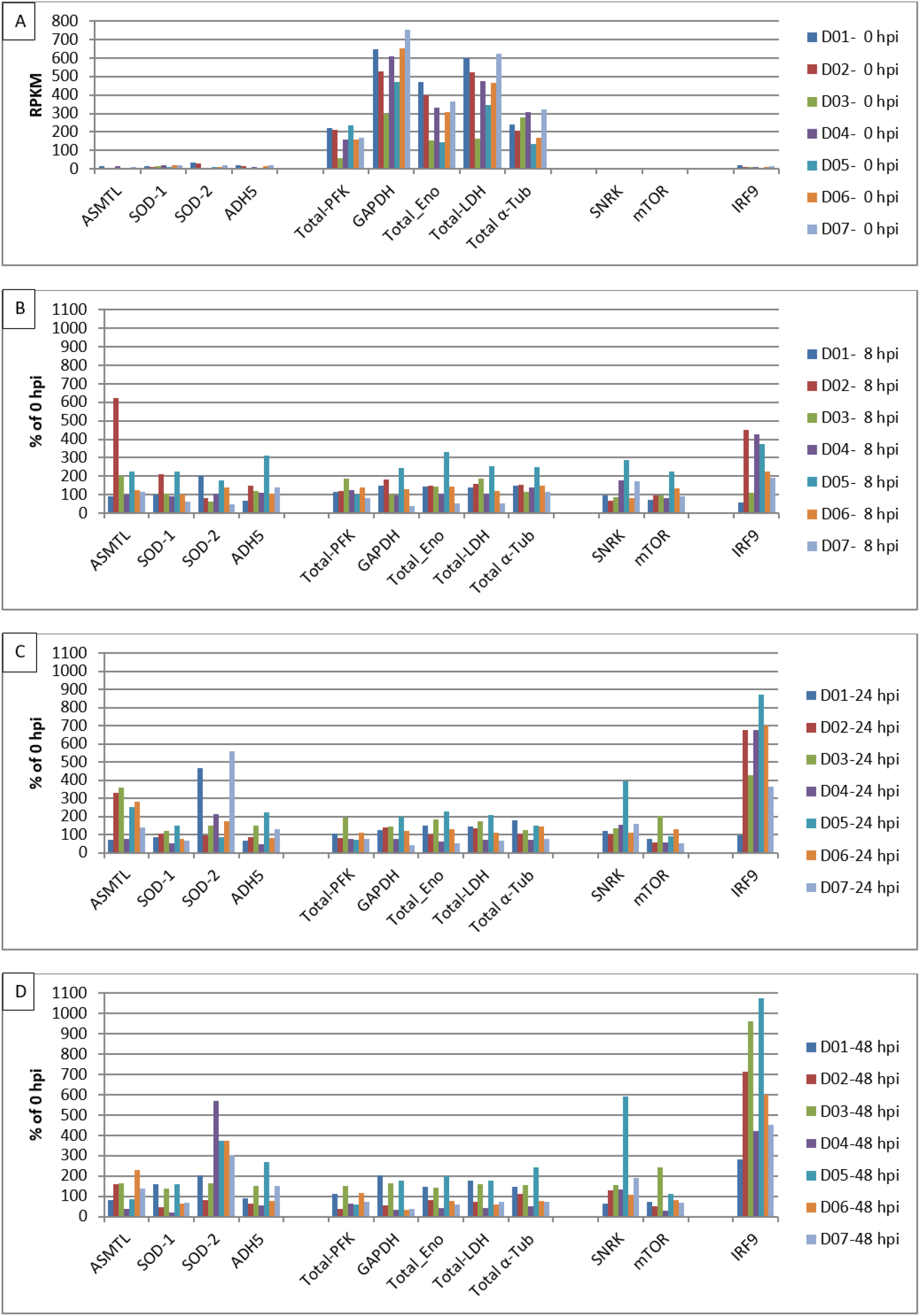
Early transcript accumulation of selected *ReprogVirus* marker genes in human nasal epithelial cells from seven donator origins infected with H3N2 Influenza virus. A: 0 hpi in RPKM; B: 8 hpi in % of 0 hpi; C: 24 hpi in % of 0 hpi; D: 48 hpi in % of 0 hpi

In **Figure 3**, transcript accumulation of eight H3N2 virus proteins is shown in RPKM for all cell donor origins at the left side to indicate virus replication/proliferation and at the right side response of IRF9 in RPKM is included. It becomes clear that IRF9 is linked in its response at transcript level to differential cell donor responses at 8 hpi and 24 hpi. This observation is supported by correlation analyses (see **Supplementary File S4**). Further, it confirms the quality of IRF9 as early responsive marker for the classical immune system. It can also be confirmed that the low response of cells from origin 1 upon influenza virus infection shown in Figure 2, was indeed associated to low virus replication. Cells from origin 1 demonstrate the lowest IRF9 response among all cell origins at early time points also when considering RPKM values. The lower values observed in general for transcript levels of influenza virus proteins at 48 hpi compared to 24 hpi and modified relations between profiles from different cell origins at that later time point seems to indicate suboptimal differential transcription rather due to non-synchronized cell cycles phases.

**Figure 3:**
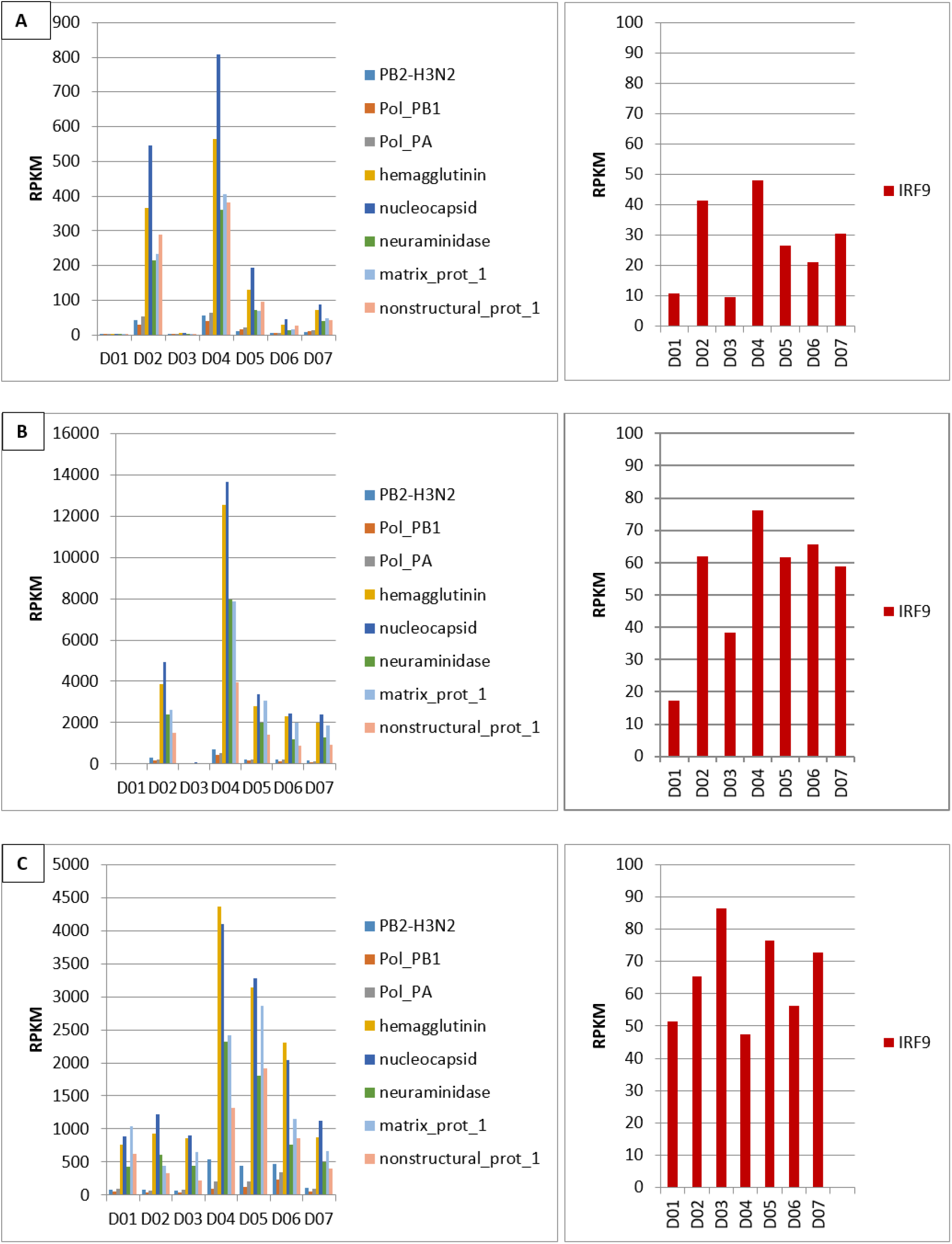
Transcript accumulation (in RPKM) of eight H3N2 virus proteins and human IRF9 in nasal epithelial cells from seven human donator origins shown at 8 hpi (A), 24 hpi (B) and 48 hpi (C).

**Figure 4** highlights the relationship between LDH transcript levels and the immune response represented by IRF9 in % of 0 hpi in human NECs for the seven cell donator origins. It can be noticed that cells from origin 1 responded in an exceptional way concerning LDH transcription and this connects to the lowest response of IRF9 among all cell origins. For the majority of cell origins LDH transcript level was increased at 8 hpi and then decreased over time. However, while cell origin 1 also shows an initial increase for LDH at 8 hpi, transcript accumulation remains at that level at 24 hpi and even is enhanced at 48 hpi. On the contrary, cell origin 5 shows highest initial increase in LDH transcript levels, which connects to the highest IRF9 response at 24 hpi and 48 hpi. Highest initial LDH transcript accumulation for cell origin 5 at 8 hpi had been combined with highest α-tub transcript levels, whereas at 24 hpi cell origin 1 demonstrated highest transcript accumulation for this gene indicating structural reorganization (**Figure 2**). Overall, and considering the combination of LDH and α-tubulin transcript level dynamics these observations point to an early indicated delay in the response to H3N2 infection for cells of origin 1.

**Figure 4:**
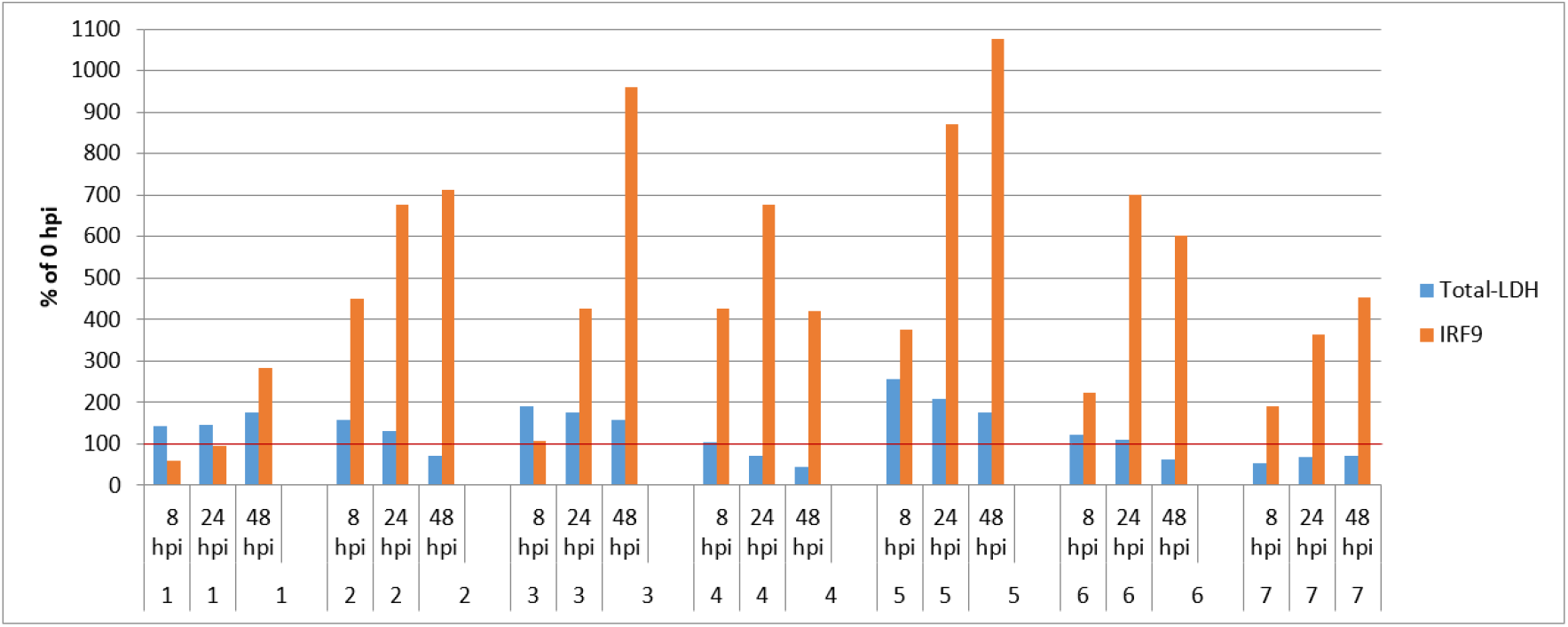
Transcript accumulation of LDH and IRF9 in human nasal epithelial cells from seven donator origins infected with H3N2 Influenza virus

## 4. Discussion

The complex marker CoV-MAC-TED was developed from research on coronaviruses using diverse cell systems (3, 4). Its relevance was recently confirmed in human NECs infected by SARS-CoV-2 variants by using a relatively small set of selected *ReprogVirus* genes (5). Here, it is for the first time that CoV-MAC-TED was studied in influenza-infected cells. Both virus types are responsible for respiratory infections with nasal cells as primary targets.

In **Figure 5**, we discuss our results integrated into a simplified scheme that was developed in an analog way by Bharadwaj et al. (6) and adapted by Costa et al. (5) for its use in virus research. This scheme considers dynamic interplay between virus infection, ROS/RNS signaling, carbohydrate stress metabolism, aerobic fermentation and cyt respiration based on knowledge related to recently published insights (3, 5) and state-of-the-art knowledge on general stress biology and virus infections. The figure is focused on the five main components of CoV-MAC-TED (a) ROS/RNS balance shift, represented by ASMTL, ADH5 as markers, (b) glycolysis and fermentation, represented by enolase and LDH, (c) structural reorganization and cell cycle progress and arrest, represented by E2F1, mTOR and α-tubulin, (d) energy status signaling, represented by SNRK and mTOR, and (e) initiation of the classical immune system, represented by IRF9. As a hypothesis, we assume that early oxidative stress associates with rapidly increased sucrose/glucose cell levels. These will stimulate the Cyt pathway via enhanced glycolysis, pyruvate production and increased TCA cycling in a way that the respiration chain gets overloaded by electrons. Consequently, ROS and RNS concentrations will increase. However, in this situation the Cyt pathway might temporarily become restricted due to rapidly consumed oxygen. In turn, aerobic lactic acid fermentation will be activated and TCA blocked through feedback control. Depending on stress level and the amount of sugar and duration of a situation of high-sugar level, anaerobic glycolysis can reach high turnover during this early phase of cell reprogramming and create a level of ATP production corresponding to the Warburg effect. Warburg effects are increasingly recognized as being part of normal physiology (28, 29) that enable host cells to rapidly mobilize energy for host maintenance and stress escape. In case of two SARS-CoV-2 variants, recent results indicated that rapidly available energy from lactic acid fermentation was first consumed for cell cycle progress (E2F1, mTOR and α-tubulin transcript accumulation signaling) and only from around 24 hpi energy was increasingly used for virus replication (5). However, our results on influenza H3N2 infection show a different scenario. Here, we did not encounter indicated cell cycle progression. To the contrary, low transcription of E2F1 upon influenza virus infection suggests that cell cycles are being arrested already very early. This can be traced through basal transcript levels of mTOR at 8 hpi and reduced mTOR transcript accumulation at 24 hpi (86%) and at 72 hpi (81%). These functional marker-based observations are in good agreement with early encountered transcription of influenza virus proteins from 8 hpi (**Figure 3**), which indicate early virus replication. Virus proliferation is accompanied by SNRK-indicated energy depletion from 8 hpi

**Figure 5:**
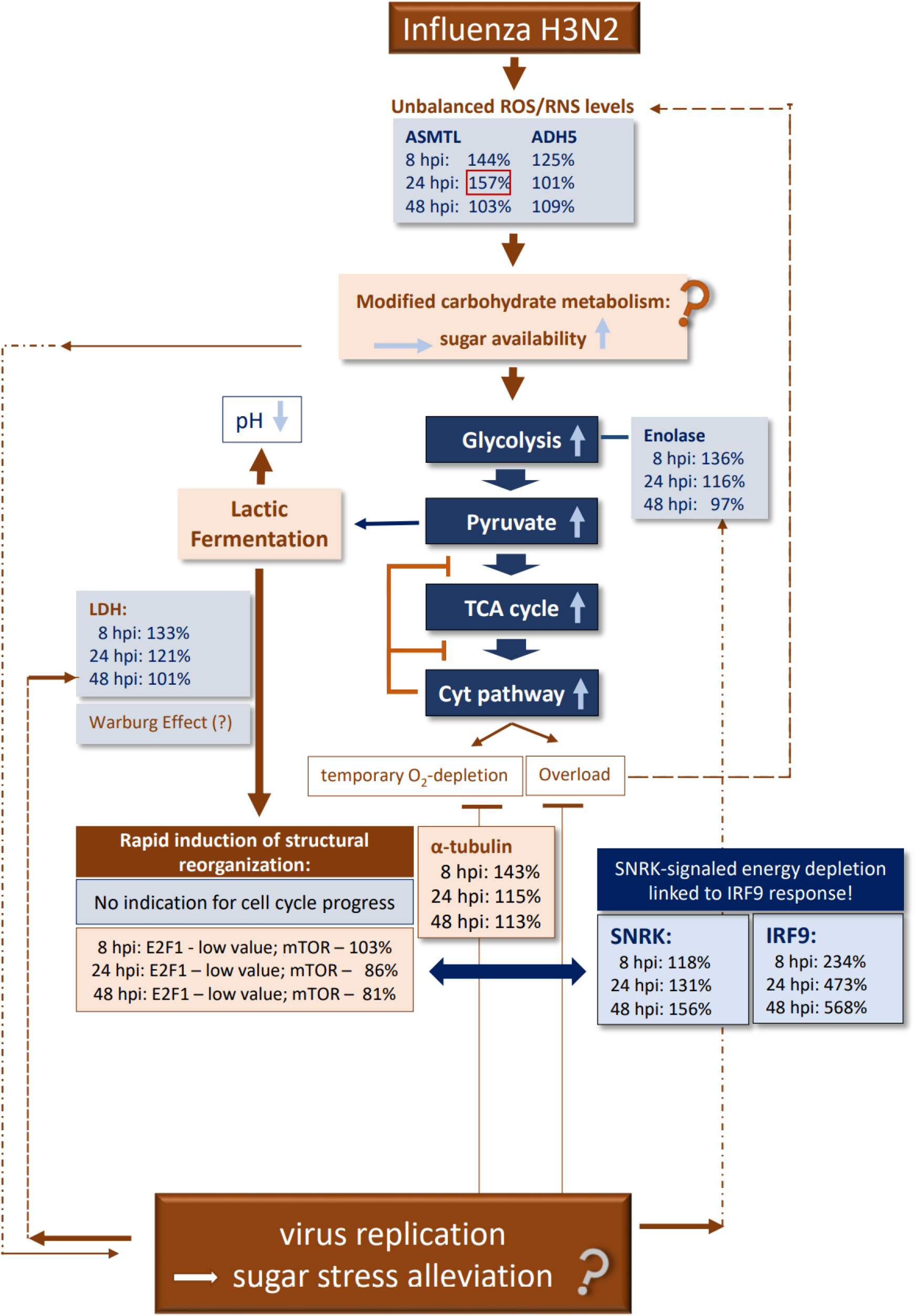
Validating CoV-MAC-TED as crucial trait for early Influenza H3N2-induced reprogramming in human nasal epithelial cells – A simplified scheme for hypothetical metabolic principles and validation of markers (118% to 156% at 48 hpi). Furthermore, SNRK-marked energy depletion is strongly associated to increasing IRF9 transcript levels from 8 hpi (234% to 568% at IRF9), which signals rapid initiation of the classical immune system. This stands in contrast to our observation for SARS-CoV-2 infection by two variants (5). There we observed a slow start of virus proliferation signals at 24 hpi and only slightly increased transcript levels of IRF9 until 72 hpi. Accordingly, no SNRK-signal was observed at 72 hpi upon SARS-CoV-2 infection time for energy depletion (76%). However, during the early phases at 8 hpi and 24 hpi post SARS-CoV-2 infection, when cell cycle progress was indicated, SNRK transcript levels had pointed to energy depletion (125%, 131%).

Overall, the conceptual line highlighted in **Figure 5** suggests the following scenario for influenza H3N2-infected human NECs: Differential increases of ASMTL and ADH5 transcripts at 8 hpi point to early unbalanced, oxidative stress (144% vs 125%) that favors ROS signaling. This is observed in a situation when virus replication and the initiation of the classical immune system were already running at high level (IRF at 8 hpi 234%). From 24 hpi, RNS stress has turned to basal level as indicated by ADH5. However, highly enhanced ROS is indicated by ASMTL transcription (157%) at 24 hpi and this interpretation is supported by the observed high increase in SOD2 transcript accumulation (311%). This is contrary to our observation for SARS-CoV-2 in comparable infection trials, where a balanced ROS/RNS increase was observed at 8 hpi, but unbalanced ROS/RNS levels at 24 hpi were due to high NO production (5). For influenza, moderate increase in glycolysis (Eno: 136%) and LDH (133%) is observed at 8 hpi, when compared to SARS-CoV-2 infection (Eno: 267% vs LDH: 251%). This coincides with lower levels of α-tubulin transcript accumulation at 8 hpi for influenza infection (143%) than for SARS-CoV-2 infection (238%) pointing thus to an advanced stage of energy-dependent metabolism-related structural reorganization by influenza observed at 8 hpi compared to SARS-CoV-2 infection, where metabolic and structural reorganization for virus replication and proliferation was obviously delayed in comparison.

Virus replication may contribute to stress relieve by providing a sink for sugars. At 48 hpi during influenza infection, when virus replication was running at high level and the immune system was continuously activated, ASMTL and ADH5 indicated only basal oxidative ROS and RNS levels. Also, enolase and LDH signaled basal glycolysis and lactic fermentation, which indicated adapted TCA cycling and functional Cyt pathway. Furthermore, SNRK signaled high energy demand for virus proliferation and immune system activation. Overall, this supposes that influenza virus structures gained control over metabolism and contributed to regain equilibrated metabolic ‘normality’ despite newly focused high energy demands.

It is noteworthy to mention that we observed CoV-MAC-TED-related meaningful differential responses upon viral infections from cells of varying donor origins. Especially, we noticed that cells of donor 1 demonstrated delayed response upon virus infection concerning both, influenza H3N2 and SARS-CoV-2 (5). This was related to differences in LDH transcript levels and its dynamic development over time.

Preliminary results point to delayed metabolic-structural reorganization upon influenza infections in comparison to other cell origins. These observations strengthen the quality of our marker system CoV-MAC-TED and make it promising to support identifying critical traits for wider anti-viral strategies.

Our analyses were performed in RNA sequencing data opened for the public by Tan et al. (1) and Gamage et al. (2). The results achieved by Gamage et al. helped us to validate our results and, additionally, to confirm IRF9 as a good marker for the classical immune system. In the same data that we used, these authors studied differential virus release kinetics and complex immune responses for influenza H3N2 and SARS-CoV-2.

## 5. Conclusions

The complex trait CoV-MAC-TED, which was initially developed from coronavirus research, can also identify early responses for influenza H3N2. This indicates its appropriateness to assist searching for common anti-viral targets in view of therapeutic design strategies. The presented marker system in combination with cultured human NECs is promising to identify differential early host cell responses upon viral infections that relate to ROS/RNS signaling, energy-dependent metabolic-structural reorganization and cell cycle regulation from individual cell origins.

## Author Contribution

JHC & BA-S conceived the basic idea and plan the study. SA performed transcriptome analyses and helped JHC in results interpretation, analyses, and final manuscript preparation. CN performed statistical analyses and helped to discuss data interpretation. BA-S reviewed data analyses, interpretation, wrote the manuscript basic draft and coordinated final manuscript discussion and revision. All authors agreed on final manuscript submission.

## Acknowledgements

The authors are grateful to Isabel Velada from the University of Evora, Portugal, for help in figure improvement. We also acknowledge the availability of Manuela Oliveira, University of Évora, Portugal, to provide comments on the manuscript during its development.

